# Applying advanced circular statistics: magnetic orientation of green toad larvae

**DOI:** 10.64898/2026.03.11.710987

**Authors:** Paula M. Helfenbein, Rachel Muheim, Magdalena Spießberger, Stephan Burgstaller, Lukas Landler

**Affiliations:** University of Vienna, Department of Behavioral and Cognitive Biology, Faculty of Life Sciences, Vienna, Austria; BOKU University, Institute of Zoology, Department of Ecosystem Management, Climate and Biodiversity, Vienna, Austria; Lund University, Department of Biology, Lund, Sweden

**Keywords:** Amphibia, Anura, magnetic compass, y-axis orientation, circular mixed-effects models, repeated measures analysis

## Abstract

Many animals use the Earth’s magnetic field as a directional reference. Among those are amphibians, which can be trained towards magnetic directions. We used mixed-effects models to analyse a dataset exploring magnetic orientation in green toad larvae. We used control simulations to test whether the modelling approach produced inflated false-positive rates and to assess power and the accuracy of directional predictions. Traditional approaches applied to average individual bearings, namely the Rayleigh test and Hotelling’s test, did not detect significant deviations from uniformity. However, the linear mixed-effects models (LMMs) applied to sine and cosine components suggested orientation components consistent with a trained magnetic response. Model predictions suggested changing directional responses during the two-minute trial and with the larvae’s activity. The trained magnetic response became more pronounced towards the end of trials and with increased activity; the untrained magnetic response was split between NE and SW. Our study represents a first attempt to use recently developed circular adaptations of LMMs with complex random-effects structures to better capture animal orientation strategies, potentially increasing statistical power and providing more nuanced insight into orientation behaviour.

## Introduction

Many animals perceive the Earth’s magnetic field, including insects [1], birds [2], turtles [3] and amphibians [4]. However, magnetic experiments can be highly variable and in several instances data obtained from different research groups did not agree [5,6]. It has been argued that experimental set-ups should be standardised between research groups and include more extensive controls [e.g., 7]. This could alleviate replication issues, and following common protocols and agreeing on necessary controls is certainly important. However, behavioural data can be intrinsically variable, due to individual variation [8], small unintentional variations in experimental procedure [9] or even differences among animal caretakers [10]. Modern statistical tools (i.e., mixed-effects models) may provide another way to control for some of this variation, especially when individuals are retested. In a recent effort to develop such statistics for circular data, Landler et al. [11] showed that linear mixed-effects models (LMMs) can be adapted for circular data by transforming bearings into their respective x and y components and using a helper variable to describe this structure in the model. In this approach, the intercept p-value indicates whether the sample deviates significantly from circular uniformity. Importantly, such an approach could provide much more detailed insights into animal behaviour than mean bearings alone. For instance, the orientations could change through time, vary between individuals, or consist of multiple overlaid responses. To put the newly developed statistics to the test, we conducted a magnetic orientation experiment with the European green toad (*Bufotes viridis*). We trained toad larvae to a light gradient, similar to the y-axis experiments successfully performed with many amphibian species [12–15]. In the wild, amphibian larvae use the magnetic field to orient along an axis perpendicular to the shore, associated with a shallow-deep and light-dark gradient [16]. We used a repeated-measures experimental design and LMMs to analyse the orientation of green toad larvae. Our study is a first step towards changing how animal orientation data are analysed and opens new possibilities for analysing repeated-measures orientation experiments.

## Materials and Methods

### Experimental set-up

Green toad larvae were kept in five transparent 4 l plastic water tanks (length 21 cm; width 13 cm; height 15 cm) with five larvae per tank from April to the end of May 2024. The training phase lasted from 7 to 29 May, and testing was performed on 30 and 31 May 2024 (see supplementary information for details on animals and husbandry). After 31 days of development in water tanks outside the coils, the tanks were positioned at the centre of a two-axis double-wrapped Merritt four square (side-lengths of 115 cm) coil system [17]. An additional Merritt coil along the North-South axis cancelled out the Earth’s magnetic field (see Fig. S1). Each individual double-wrapped Merritt coil system was powered by a digitally controlled DC power supply (KORAD KD3005D DIGITAL-CONTROL 0-30V 0-5A). Coils were controlled by a relay board, with two relays controlling the polarity of each double-wrapped coil. The Merritt coil system was surrounded by a Faraday cage consisting of two layers of 1 mm wire mesh to block possible disturbance from radio-frequency fields [18], with no electric devices inside the cage except a light source and a camera that were mounted at the top centre of the Merritt coil system for video recording. Magnetic fields were measured with a magnetometer (Fluxgate Magnetometer Fluxmaster, Stefan Mayer Instruments GmbH C Co. KG). The tanks were illuminated by four 230 V AC LED panels (Pawel LED module 107219 230V 4000K), positioned 138 cm above the tank floors. At a height of 60 cm above the tanks a translucent glass pane ensured a homogeneous light distribution (Fig. S1, see supplementary information for light intensities).

### Training phase

For the training phase the Earth’s magnetic field was cancelled out and an artificial magnetic field with an intensity of 49.3 µT (1.0 A), matching the local Earth-strength magnetic field, was generated in the direction of magnetic North (= geographic North). One side of each tank was covered with black cardboard, creating a light-dark axis, simulating the light gradient in natural spawning waters. We defined the trained direction as the direction from the dark side towards the light side, which was oriented in a different direction for each tank relative to the generated magnetic field, creating four different training directions (Fig. 1A, Fig. S2). An air pump and food were always positioned on the light side of each tank. White paper blinds between tanks blocked the view from one tank to the next. Tanks were positioned randomly with regard to their training direction within the centre of the Merritt coil system. Two animals died three days after the training phase started and were replaced with new individuals. One animal died during the training phase before testing.

**Figure 1:**
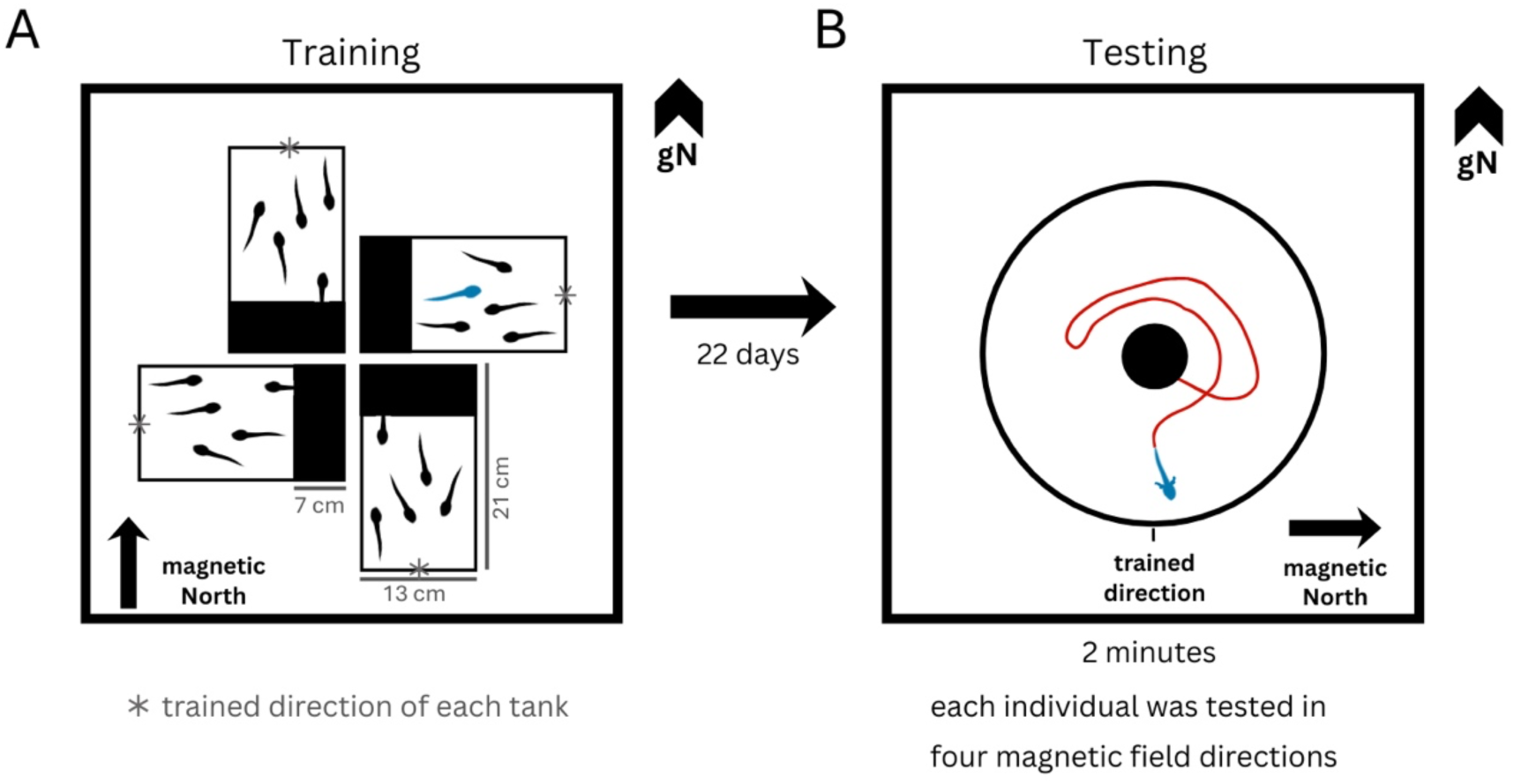
Schematic illustration of the experimental procedure. (A) Green toad larvae were placed in rectangular tanks inside the Merritt coil system, with the magnetic North (mN) in the direction of geographic North (gN). One side of each tank was covered with black cardboard to create a light-dark axis, oriented differently relative to mN for each tank. White paper was placed between tanks to block visual cues from other tanks. The air pump and food were always positioned on the light side of the tank. Following a 22-day training period, testing was conducted in the same Merritt coil system. (B) Training tanks were replaced with a circular, water-filled testing arena (diameter: 37.5 cm). Each larva was individually transferred to a release device located at the centre of the testing arena. Each larva was tested four times, with mN aligned to a different geographic direction in each trial.

### Testing procedure

After a training phase of 22 days twelve tadpoles (median Gosner stage: 37, [19], see supplementary information) were tested. Immediately before testing the animals were separated into individual closed rectangular plastic containers (length 18 cm; width 6.5 cm; height 8 cm) with 3 cm water depth, in the dark. During testing, these containers were positioned outside the Merritt coil system and were therefore exposed to the natural magnetic field, but with the axis of the containers always aligned with their trained direction. Testing was performed within the same set-up as the training, with the same Merritt coil system, light setup, and Faraday cage (Fig. S1B). We created a magnetic field in the required testing direction with an intensity of 49.5 µT (± 0.7 µT) (1.0 A) by only changing the polarity of single coils in the double-wrapped system, without turning any coils off. After a 5-minute settling period in the individual containers, tadpoles were individually transferred to the testing arena and released using a release device after a 10-second acclimation (see Fig. S3 and supplementary details). The experimenter operated the release device from the geographically southern side of the setup. Each tadpole was tested four times, each time with the magnetic field in a different geographic direction (Fig. 1B), but representing one experimental treatment, as the setup was otherwise identical. Between tests, they were returned to their individual containers. The order in which the larvae and magnetic field orientations were tested was randomised, therefore the time between trials for each individual varied, with an average of 36 (SD: ± 34) minutes and a minimum of 5 minutes. To avoid directional chemical cues, the release device was rinsed with tap water after every test and the arena water was stirred before each test. Tadpole behaviour was recorded using a camera (ELP camera ELP-USB16MP01-BL170) mounted centrally above the testing arena. Nine of the 48 trials were excluded because of release errors: the release device did not sink down completely, the larva escaped before the device was fully opened, or the larva did not leave the device within the trial period. After exclusions, the dataset contained 39 valid trials from 11 individuals.

### Tracking

We used a custom-made tracking software written in MATLAB (Version R2023b), MathWorks, to extract the central position of each tadpole at 15 frames per second (see supplement for video preparation and tracking details, videos can be accessed online [20]). The positions of the tadpoles in the arena were calculated relative to the centre of the arena to obtain the direction and an r-value (distance to centre when the radius equals one) for each position. As a measure of activity, we calculated the distance moved by the tadpole between frames relative to the radius of the arena.

### Statistical analysis and plotting

All analyses were performed in R [21] (see the supplementary material for details on packages, data files and R code). We used the recently published mixed-effects approach for circular data [11]. In this approach, the x and y components of the directions [cos(θ), sin(θ)] are used as the response variable. The distinction between x and y (cosine versus sine) is coded using a helper variable (cosine coded as the base level). We calculated four different samples: orientation relative to geographic North (gN, ignoring the trained directions and magnetic field), orientation relative to magnetic North (mN, ignoring the trained directions and geographic North), orientation relative to a potential trained geographic direction (gT, ignoring the magnetic field and an untrained geographic orientation), and orientation relative to the trained magnetic direction (mT, ignoring the geographic North and an untrained magnetic response) (Fig. S2). Animals were trained in one of the four cardinal magnetic (and geographic) directions and tested in each of the four cardinal magnetic directions, to be able to differentiate between geographic and magnetic directional preferences. The same models were applied to all four scenarios. We assumed that the larvae’s vector length (“r”-value) and activity, both in interaction with time during the trial (zero to two minutes), influenced orientation, and therefore added these factors to the model. The vector length was shifted so that its lowest observed value was zero (r_t = r − min(r)), so that the intercept refers to the lowest observed vector length rather than to r = 0, where no direction is defined. The random-effects structure of our model included a random intercept for ID and random slopes for both time and test round within ID (random structure: “Time + Test_round | ID”). We detected no multicollinearity issues. Note, although we performed several simulations to evaluate the approach (see Control simulations), confidence intervals and p-values should be treated with caution, as this approach is still experimental. In addition, we applied the traditional Rayleigh test on mean angles for each individual and the Hotelling’s test on the individual weighted mean vectors.

### Control simulations

The approach is still novel and clearly violates the assumption of normality (cosine and sine have finite limits [-1,1], see Fig. S4) and certain situations are still untested (e.g., random slopes); therefore, we performed pilot simulations to test potential type I error issues, the usability of the (intercept) p-value, effects of different ways to include the helper variable in the model, and the accuracy of the predicted directions. In brief, we fitted our full models to uniform samples to estimate the type I error rates of all model terms (Table S6), and we analysed samples drawn from von Mises distributions with different mean directions and concentrations with an intercept-only model to estimate the power of the intercept p-value and the accuracy of the predicted directions (Figs S6, S7). We further simulated a more complex scenario with an interaction between time and activity to compare different ways of including the helper variable in the model (Fig. S8), and added repeated testing of individuals with individual-specific deviations to compare different random-effects structures (Fig. S9). In addition, we performed a limited sensitivity analysis by thinning the real data of the trained magnetic response (Fig. S5). Details of all simulations are given in the supplementary material.

## Results

Neither the Rayleigh test nor the Hotelling’s test detected significant deviations from uniformity in any sample (Fig. 2: A, B, E, F, I, J, M, N). Taken together, the LMM predictions indicated directional responses in the gN, mN, gT and mT reference frames (Fig. 2 and 3; Tables S1-S4), however the mT response showed a clear predicted unimodal orientation towards the expected direction. Non-uniformity in multiple reference systems is plausible for at least two reasons: 1) the larvae could exhibit multiple different behaviours during the trial (e.g., orienting away from the experimenter at first and searching for an escape using magnetic cues later in the trial); 2) our data set was not completely balanced (some trials were unsuccessful), therefore strong directionality in one reference system could create directional biases in another. The geographic orientation was directed towards North (away from the experimenter who started the trial) at the start of the test, shifted westwards during the trial, and was almost unaffected by activity (Fig. 3). The magnetically trained orientation started with a westerly biased orientation and shifted during the trial towards the expected direction, and exhibited less variance during high activity, as shown by its narrower confidence interval (Fig. 3). The trained geographic orientation resembled the magnetically trained orientation, except at high activity (Fig. 3). An untrained magnetic response towards NE was most pronounced at the start of the trials, while activity was associated with a switch from NE (low activity) to SW (high activity) (Fig. 3).

**Figure 2:**
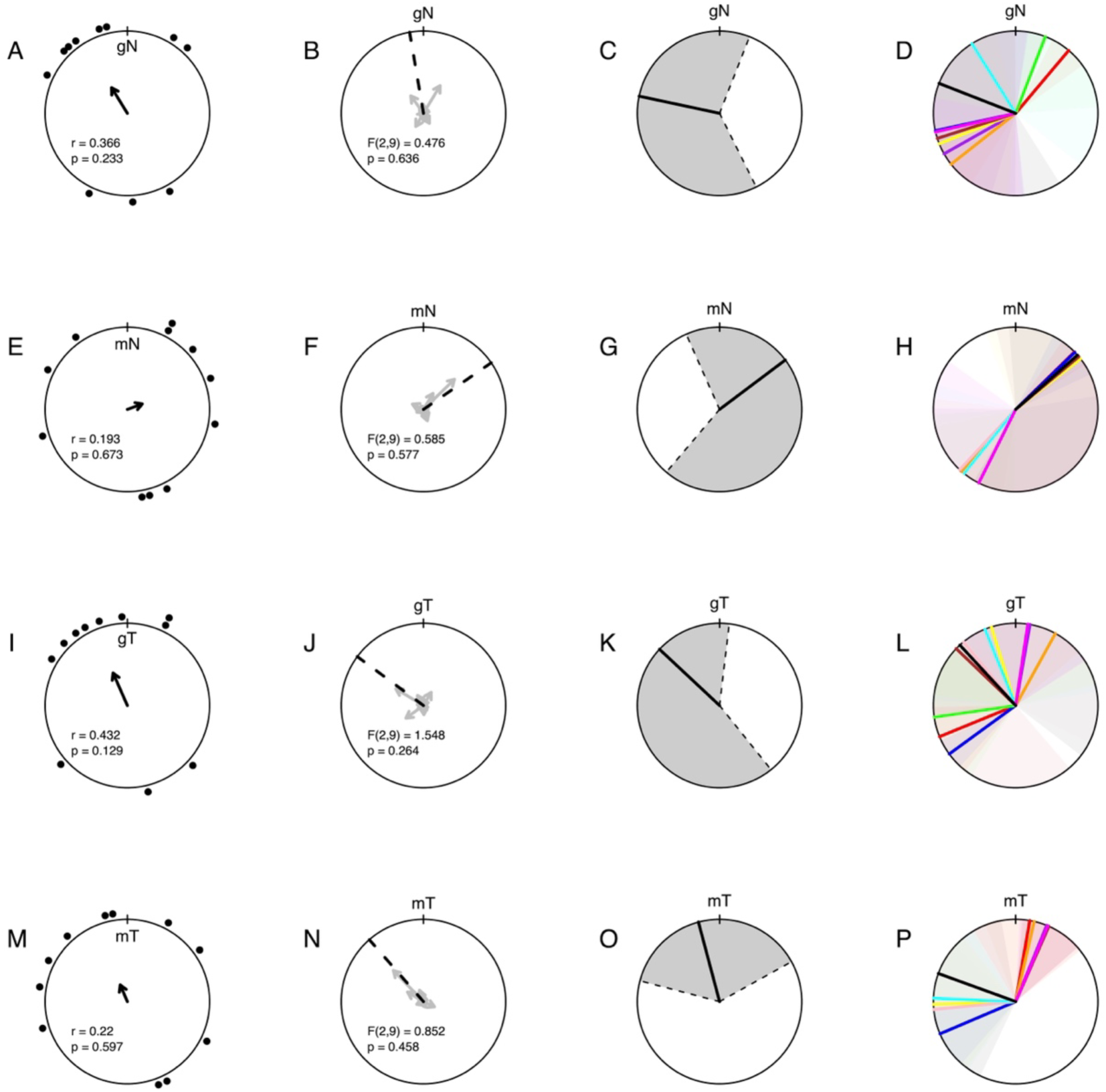
Results of the magnetic orientation tests of green toad larvae using four different approaches and visualisations. Data are shown with respect to the geographic North (gN, A-D), magnetic North (mN, E-H), the geographic trained direction (gT, I-L), and the magnetic trained direction (mT, M-P). The first order responses (circular mean bearing of each individual, disregarding vector length) were tested against uniformity using the Rayleigh test (A, E, I, M), r and p-values for each of the samples are shown in the plots. The mean vector of each sample is indicated with a black arrow (circle radius indicates r = 1). The second order responses (weighted circular means for each individual, taking the vector lengths into account) were tested against uniformity using the Hotelling’s test (B, F, J, N), the F (degrees of freedom in parentheses) and p-values are shown inside the plots. Mean individual vectors are shown in light grey, the mean direction is shown as a dashed black line (circle radius indicates a vector length of one). The average predictions (all independent variables held at their average values) were based on the intercept result of the LMMs (C, G, K, O). We added grey shaded 95% confidence intervals (CI) to the mean predicted direction (black line). In addition, the individual predictions are shown (D, H, L, P), each individual has the same colour in all samples, CIs are shown for all individuals to exemplify the data structure and spread, in the same colour as the predictions but slightly translucent to allow overlaying colours. The individual predictions (including CIs) are listed in Table S5.

**Figure 3:**
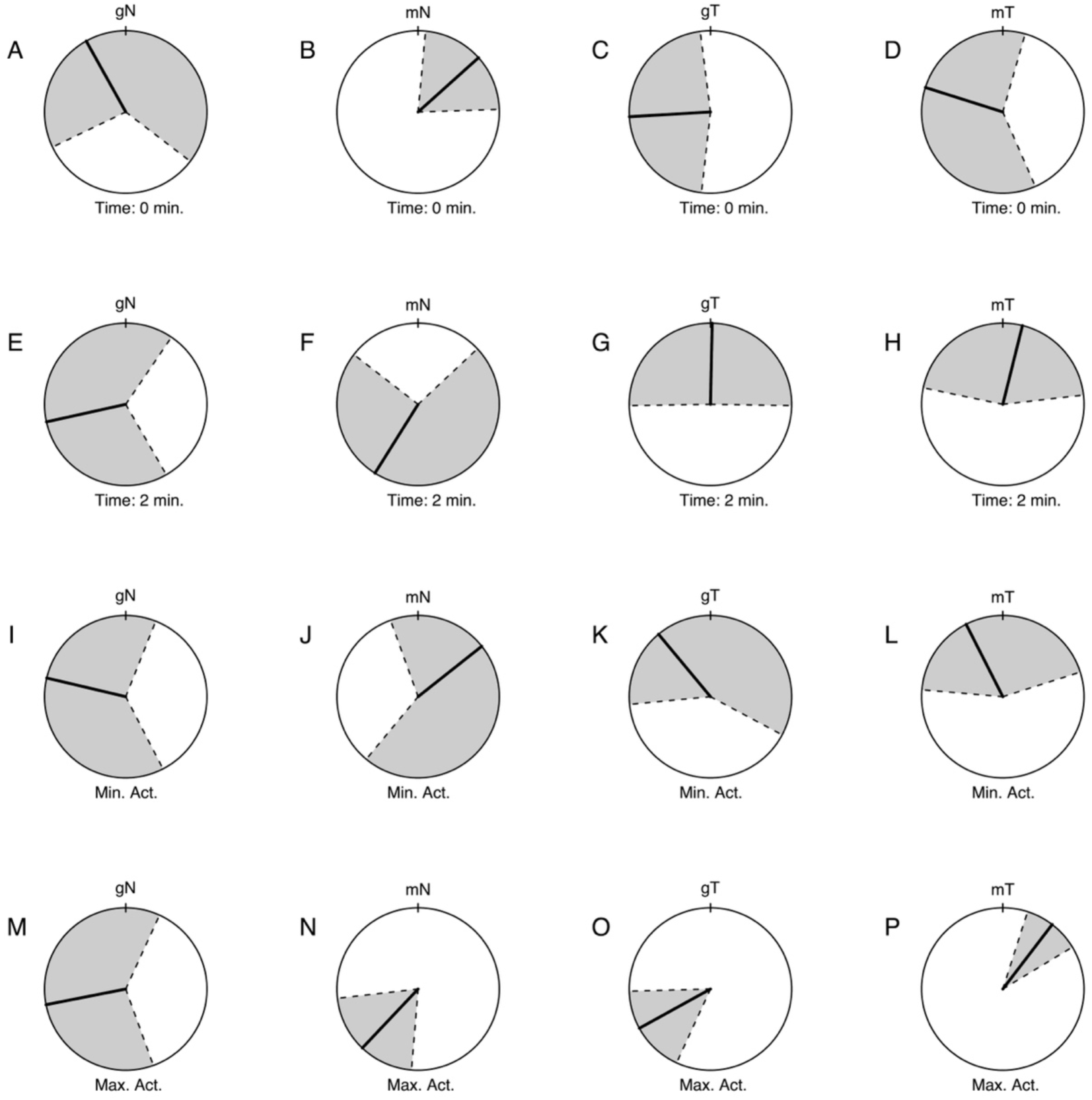
Predictions based on the LMM models using the minimum and maximum values of time (zero minutes (A-D) and two minutes (E-H)) and activity (minimum (Min. Act., I-L) and maximum activity (Max. Act., M-P), while keeping the other independent variables at their averages. Samples are analysed with regard to geographic North (gN, A, E, I, M), magnetic North (mN, B, F, J, N), towards the geographic trained direction (gT, C, G, K, O) as well as towards the magnetically trained direction (mT, D, H, L, P). Mean predictions (black line) are shown with their 95% confidence intervals (grey shaded area).

On the individual level orientations with regard to gN, gT and mT all appeared to be anticlockwise biased (Fig. 2: D, L, P), while orientations with regard to mN showed a clockwise bias (Fig. 2: H). However, there was an evident similarity between gT and mT with an approximate 90° split between two groups of individuals, one towards the expected direction and one anticlockwise. The individual predictions in relation to mN appear to be split between two highly clustered opposite groups (around 45° and 220°), potentially indicating a bimodal distribution, which we did not evaluate further.

Our control simulations yielded type I error rates close to the nominal level for all model terms and in all simulated scenarios (Table S6; Figs S7, S8 and S9). The sensitivity analysis indicated stable model behaviour with real data, as the predicted mean direction for mT did not change when the data were thinned (Fig. S5). Predictions from uniform and axial samples were highly variable and therefore not usable, whereas predictions for unimodal samples were at the expected direction, with decreasing variability as the concentration increased (Fig. S7). In the more complex scenario with an interaction between time and activity, predictions remained close to the expected direction irrespective of how the helper variable was included in the model (Fig. S8). Adding individual-specific errors increased the variability of the predictions, but all random-effects structures yielded nearly identical predictions (Fig. S9). In contrast, with the cosine as the base level of the helper variable, the p-value of the intercept only had usable power to detect clustering towards or opposite to the direction represented by the cosine component. For clustering perpendicular to this axis and for axial samples, its power was low, also in the more complex scenarios (Figs S6, S8 and S9).

## Discussion

We show that our novel statistical approach - i.e., incorporating mixed-effects model structure in circular analyses - can provide a flexible model-based analysis of magnetic orientation experiments and even shed light on fine-scale structure in individual behaviour. Although only eleven animals were tested, our model-based analysis provides evidence consistent with magnetic orientation in green toad larvae. Our control simulations and sensitivity analysis indicated that the approach does not produce inflated false-positive rates under the simulated uniform-response scenarios, irrespective of how the helper variable or individual random effects were included, and is robust to data thinning. Our simulations also revealed an important limitation of the intercept p-value: with the cosine as the base level of the helper variable, it only detects clustering towards or opposite to the direction represented by the cosine component, whereas clustering perpendicular to it and axial distributions remain largely undetected. In contrast, predicted directions were accurate for unimodal samples, also in the more complex scenarios and regardless of how the helper variable was included in the model. The choice of random-effects structure for the helper variable did not affect the predictions.

Interestingly, none of the traditional tests detected deviations from uniformity, potentially indicating that the model-based approach can detect structure that is lost when repeated observations are reduced to individual mean bearings. The orientation characteristics differed intriguingly among the reference frames. At this stage, detailed interpretation of these characteristics remains speculative. However, the geographic orientation (stronger at trial start) might be consistent with an escape response away from the experimenter starting the tests. The magnetic orientation (disregarding the trained direction) may indicate an orientation along the NE-SW axis, shifted clockwise relative to magnetic North. Such orientation has been shown in many magnetic behavioural experiments without directional training or when animals did not follow the trained direction [22]. Such untrained (spontaneous) magnetic preferences have been interpreted as a possible behaviour to integrate multiple cues in a reference frame [23–25] and also in this experiment it was most prominent at the trial start. However, spontaneous magnetic behaviour remains poorly understood and warrants further experiments and analyses. Both trained responses (in relation to gT and mT) showed a possible 90° split between two groups of individuals, although only the magnetically trained response showed an overall directionality towards the trained direction. In magnetic experiments with mice such a split around the trained direction has been shown when trained towards magnetic East [26]. It has been argued that this split may originate at the receptor level [27]. However, in our case, the split was independent of training direction and present also in the geographic response. One possible explanation is edge-following behaviour (similar to the behaviour of mice in an open field [28]): larvae trained in rectangular containers may have associated the trained axis with movement towards the combination of the light-dark axis and the nearby wall. Such a conflict could shift the observed direction away from the trained axis and potentially split the animals into two groups. The anticlockwise bias in both trained directions (gT and mT) may reflect left-right asymmetries, which needs further exploration. In conclusion, this pilot dataset illustrates that circular adaptations of mixed-effects models can provide detailed, model-based insight into repeated-measures orientation data. The approach detected orientation structure that was not apparent from traditional analyses of individual mean bearings, and our simulations indicate accurate directional predictions for unimodal responses. However, further work is needed to develop mixed-effects models for circular data, to evaluate the approach for axial responses, and to assess its performance across a wider range of experimental designs.

## Supporting information

Data and Scripts

## Acknowledgments

We thank Daniel Grötike, Norbert Schuller and Astrid Pledermann for their help constructing the coil setup, Christian Gützer for his help with the control system of the coils, Ethan Stewart for designing, 3D-printing, visualising the release device, and three anonymous reviewers for their highly valuable comments.

## Ethics

Animal studies were performed in accordance with the Austrian federal ministry (2020-0.814.136) and the City of Vienna (MA22-230917-2020).

## Data availability

All data files and R code can be found within the article and its supplementary information. Videos have been uploaded to Zenodo [20].

## Supplementary Methods

### Experimental animals and husbandry details

Spawn of the green toad (*Bufotes viridis*) was collected from the Rudolf-Bednar-Park in Vienna (48.2260° N, 16.397° E) on 6 April 2024 from an artificial pond. Training and testing took place in the Gregor-Mendel-Haus of the BOKU University in Vienna (48.2365° N, 16.3370° E). Larvae were kept in five transparent 4 l plastic water tanks (length 21 cm; width 13 cm; height 15 cm) with five larvae per tank. Each water tank was filled with tap water and equipped with an air pump for oxygen supply. All tanks were kept under the same environmental conditions following the outdoor temperature with a 12-hour light-dark cycle. Larvae were fed three times a week with 0.5 g of fish flakes (Futter-Flakes Flockenfutter für Zierfische, Centra24 e.K., Germany) per animal and once a week with one fish pellet (JPL NovoPleco 30310 Complete Food for Suction Catfish Tablets, JBL GmbHCCo.KG, Germany).

**Figure S1:**
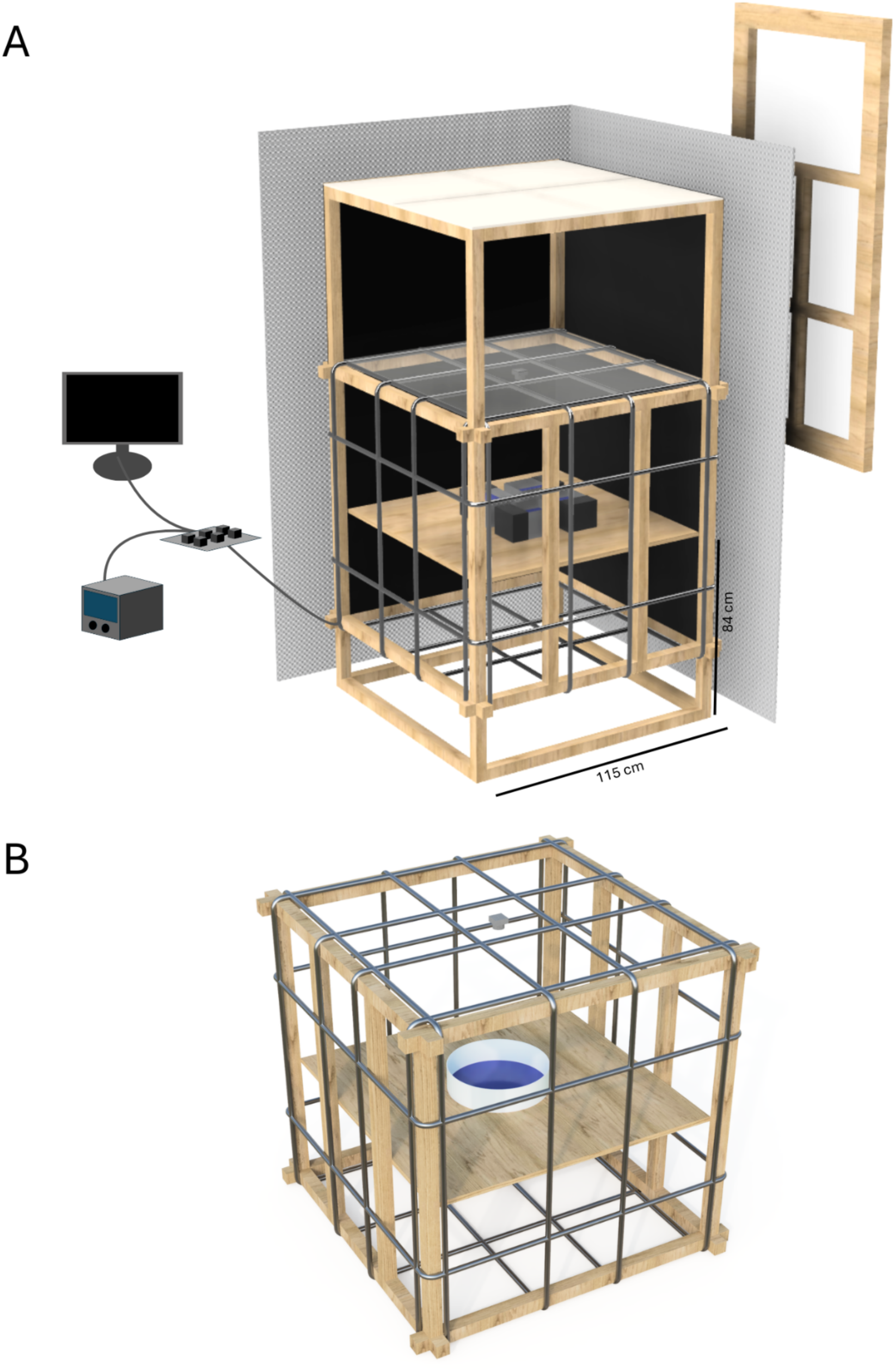
Schematic of the complete experiment setup. (A) At the top, four LED panels illuminated the setup, with a translucent glass pane beneath them to homogenize the light distribution. A two-axis Merritt four-square coil system with double-wrapped coils generated a horizontal magnetic field. An additional Merritt coil system along the North-South axis allowed us to cancel out the Earth’s magnetic field. A table was placed in the centre of the Merritt coil system on which the training tanks or the testing arena could be placed. At the top a camera was mounted for video recording. The setup was covered with black light-proof curtains on all sides and placed inside a Faraday cage consisting of 1 mm wire mesh. All coils were powered by a power supply and controlled by a relay board and a computer, set up outside the Faraday cage, approximately one metre away. Towards geographic East of the setup, behind the curtain and the Faraday cage, was a closed window. For the training phase tadpoles were placed in training tanks with one side always covered with black cardboard. (B) For testing the training tanks were replaced with a testing arena using the same magnetic coil setup.

### Light intensity measurements

The light intensity at tank height was 60 x 10^17^ quanta s^-1^ m^-2^ (±6 x 10^17^ quanta s^-1^ m^-2^) with a spectral distribution of 9.7 x 10^17^ quanta s^-1^ m^-2^ blue light (400-500 nm), 28.1 x 10^17^ quanta s^-1^ m^-2^ green light (500-600 nm), 23.4 x 10^17^ quanta s^-1^ m^-2^ red light (600-700 nm) and 2.1 x 10^17^ quanta s^-1^ m^-2^ above 700 nm. The setup was shielded from other light sources by a black curtain on all sides. Light intensities were measured with a UPRtek Spectrometer (PG100N Spectral Par Meter) in µmol s^-1^ m^-2^ and converted into quanta s^-1^m^-2^.

### Gosner stages

Twelve individuals were tested with Gosner stages ranging from 27 to 42, with a median Gosner stage of 37. Animals above Gosner stage 44 were excluded (12 animals) because they had progressed too far in their metamorphosis, hence, they had developed all four limbs and could not be tested in our release device.

### Details for experimental setup

**Figure S2:**
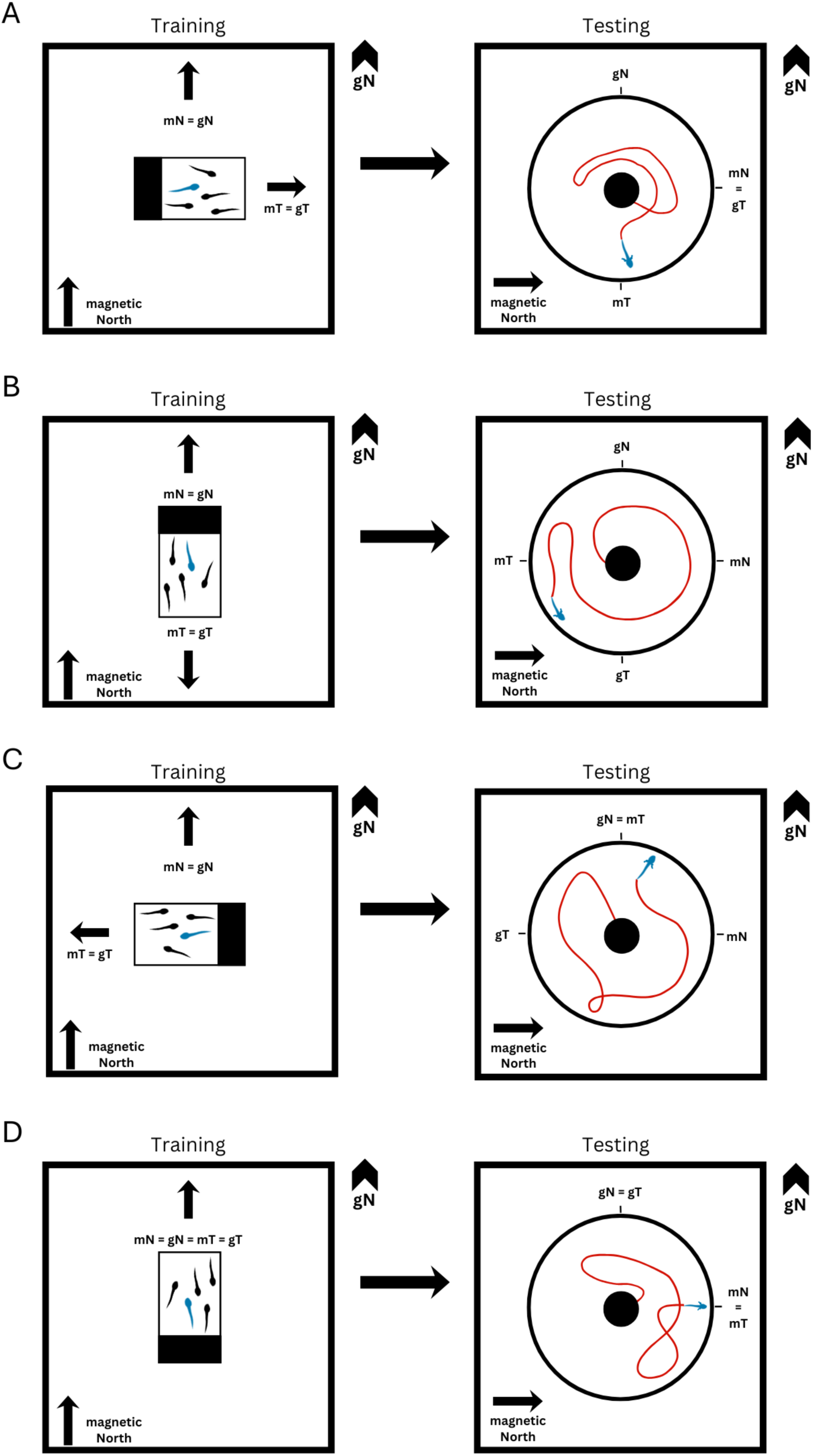
Schematic illustration of the training directions of each tank. Vectors indicate magnetic North (mN), geographic North (gN), the magnetic trained direction (mT) and the geographic trained direction (gT). mT and gT depend on the orientation of the training tank towards East (A), South (B), West (C) or North (D) during training. For testing a magnetic field aligned towards geographic East is shown for illustrative purposes, but for each trial for each larva the magnetic field was aligned to a different cardinal direction (East, South, West and North).

#### Details on the test arena and release device

The test arena was a round white container (37.5 cm in diameter; height 9.5 cm) filled with tap water and 10 ml of water from the original training tank (to generate a familiar environment) with a water depth of 7 cm. Water temperature was maintained at 22°C (± 1°C) during the testing phase. Tadpoles were placed into a release device at the centre of the testing arena through an overlapping opening in the light curtain. The tadpole remained in the motionless cylindrical release device (4.5 cm diameter; 2.5 cm above the arena floor) for 10 seconds to settle again before the inner part of the release device, with the tadpole inside, was released by the experimenter and sank down 4 cm in 14 seconds (± 1.4 seconds), allowing the tadpole to be released underneath the water surface through 12 identical doors (1 cm height, 0.8 cm width) arranged radially around the release device (Fig. S3, Movie S1).

**Figure S3:**
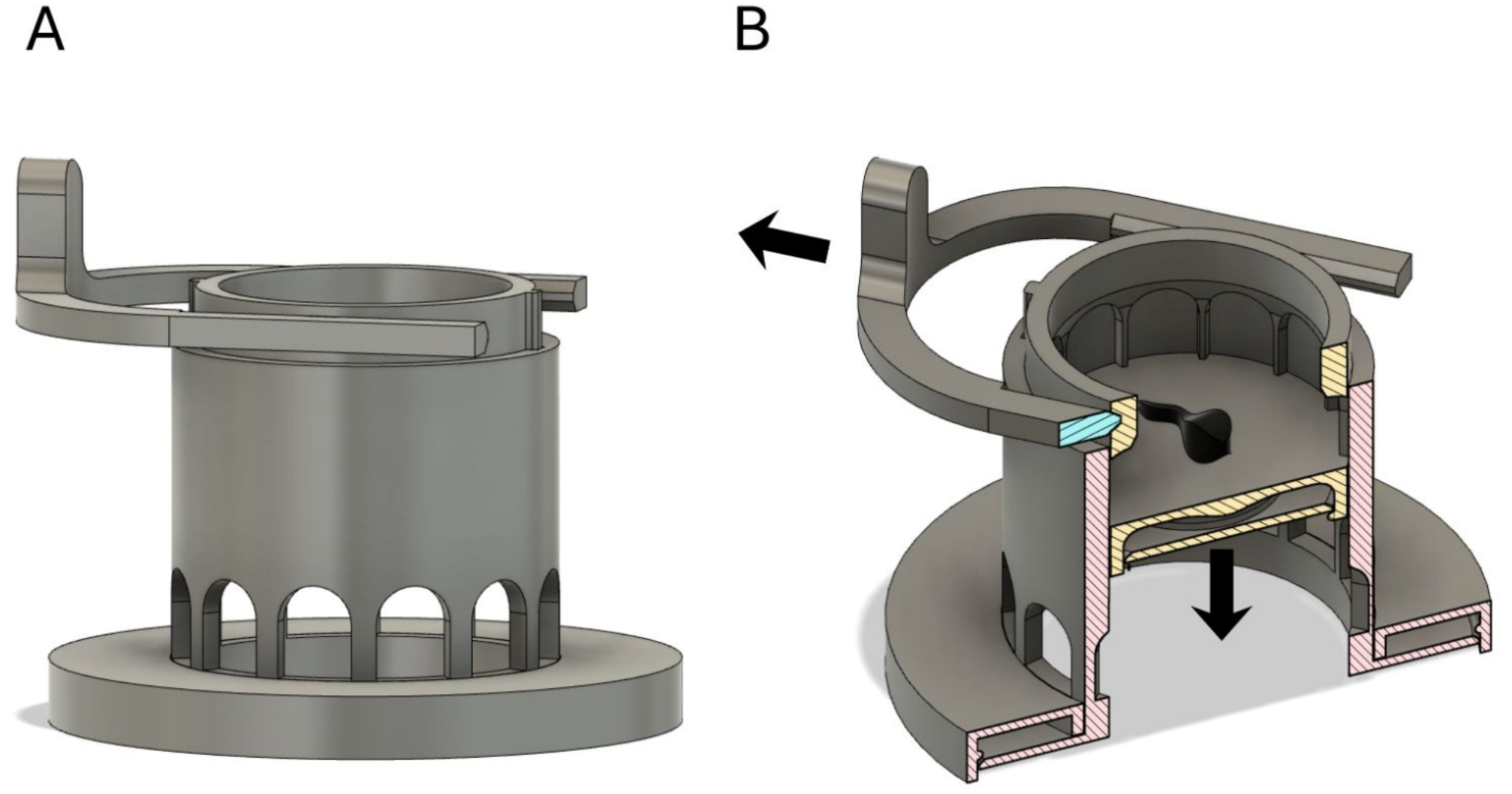
Schematic of the release device used during testing. Larvae were placed in the motionless cylindrical inner part of the release device (4.5 cm diameter; 2.5 cm above the arena floor). After a settling period of 10 seconds the experimenter released the inner part (containing the larvae) by pulling the handle to one side. This allowed the inner part to fill with water and sink down 4 cm inside the cylindric outer part of the release device in 14 seconds (± 1.4 seconds). Once the inner part was at the bottom, 12 radially arranged doors (1 cm height, 0.8 cm width) in the outer part aligned with 12 identical doors in the inner part and therefore allowed the larvae to be released through one of the doors. (A) Side view of the release device in its initial position with the handle still on, to keep the inner part fixed in the upper position. (B) Cross-section through the release device showing the inner part (yellow cut) inside the outer part (red cut) and the handle (blue cut) fixing the inner part in its initial position, with a larva inside the inner part. Arrows indicate movement of handle and inner part when released.

#### mp4 file

Movie S1: Animation showing the release device in operation. The handle is pulled to the side and releases the inner part to move inside the outer part to the bottom, therefore allowing the larva to be released through one of the doors.

### Video preparation

Videos were taken at 15 frames per second and a resolution of 1920 x 1080 and were cut to 120 seconds from tadpole release. The videos were then rotated so that geographic North was always at the top. Video files were processed using Any Video Converter (Version 9.0.1, Anvsoft Inc.).

### Tracking details

In the tracking analysis each frame was compared to the first frame of the respective video to detect changes in position and movement. For consistent spatial alignment across sessions, a mask corresponding to the upper border of the behavioural arena was created separately for each of the two testing days. This ensured precise spatial alignment and was then applied uniformly across all trials conducted on the respective day. Tracking results of every video were checked manually afterwards, and in case artefacts other than the tadpole (e.g., water movement) were tracked, these were manually removed in the tracking software.

### Statistical details

We defined significance as a p-value of < 0.05 throughout. For the linear mixed effects models the function *lmer* (packages *lme4* [1] and *lmerTest* [2]) was used, however, because of the novelty of the method we removed all mention of p-value significance for the linear mixed effects models from the main text. Potential multicollinearity was tested using the function *check_collinearity* from the package *performance* [3]. We used the *predict_response* function (package *ggeffects* [4]) to predict the mean responses and 95% confidence intervals for each model used. The x and y directional components were back-transformed to two-dimensional directions (angles) for visualisation and data interpretation. For plotting we used a customised version of the function *plot.circular* (package *circular*lin [5]). In the control simulation of the full model we generated uniform random directions using the function *rcircularuniform* in package *circular*. For the simulations with the intercept-only model, samples were drawn from von Mises distributions using the function *rvonmises* from the package *Rfast* [6], and these simulations were run in parallel using the packages *foreach* [7] and *doParallel* [8]. For the more complex simulations, samples and individual-specific deviations were drawn using the function *rvonmises* from the package *circular*. From the same package we used the function *rayleigh.test* to test against a uniform distribution for the individual mean angles (using *mean.circular* and ignoring r-value). We applied a one-sample Hotelling’s test (*T2.test* from the package *rrcov* [9]) to the individual (second order) mean vectors (calculated using *weighted.mean.circular,* taking the r-values into account) to test against a uniform distribution.

### Control simulations

For all simulations we drew 1000 independent samples from the distribution in question. First, we tested whether p-values below 0.05 could arise (using our model structure) when including random directional samples. Therefore, we drew uniform samples and used them as circular response variables in the otherwise identical models used in this paper (same independent data, same random error structure). We extracted the p-values of all six model terms (intercept, time, r, activity, time × r, time × activity) and calculated the type I error rate as the number of p-values below 0.05 (the alpha level) divided by 1000 (the number of samples), the expected type I error was therefore 0.05. Using our full model for the trained magnetic response (mT) and the real data, we further performed a limited model sensitivity analysis by thinning the data and comparing results (Fig. S5). We then drew samples (n = 1000) from von Mises distributions with κ = 0, 0.5, 1 and 2 (unimodal means: 0°, 90°, 180° and 270°; bimodal means: 0°/180° and 90°/270°, with half of each sample per mode). We analysed them with an intercept-only linear model (response ∼ helper variable). Power was calculated as the proportion of samples with an intercept p-value below 0.05. To evaluate the directional predictions, we back-transformed the predicted intercept into an angle and calculated its deviation from the expected direction (for bimodal samples, angles were doubled before this calculation). Deviations were summarised as circular mean ± circular SD across the 1000 samples. Next, we generated samples following a more complex scheme, with two added main effects, time and activity (values from 0-1), which corresponded to a directional change (0-180°) or change in clustering (κ = 0-2). In the interaction scenario, the mean direction changed with activity from 0° to 180°, and the concentration increased with time and with the sum of time and activity (κ = 2·time + (time + activity)/2, range 0-3). As the mean direction varied between 0° and 180°, we defined 90° as the expected direction. In this analysis we also evaluated possible effects of including the helper variable as a main effect only (variable + time × activity), only in interaction with the main effects (variable:(time × activity) + time × activity), or as both main effect and interaction (variable × (time × activity)); all three models were fitted as linear models. Furthermore, we added simulated repeated testing of individuals with either a bias of -20° or +20°, and kept their individual bias throughout retesting. We used the same main effects structure for all models (variable × (time × activity)) but varied the random-effects structure: (1 | ID), (1 | ID:variable), (1 + variable | ID) and (1 + variable || ID), exploring possible effects of including the helper variable.

**Table S1:** Model results table of linear mixed effects model for circular data, using the geographic response for the entire test duration.

| **Predictor** | **Estimate** | **SE** | **df** | **t** | **p** |
|---|---|---|---|---|---|
| Intercept | 0.514 | 0.04 | 12.239 | 12.7 | <0.001 |
| Helper variable | -0.047 | 0.004 | 139367 | -12.795 | <0.001 |
| Minute | -0.83 | 0.058 | 13.286 | -14.324 | <0.001 |
| r_t | -0.846 | 0.028 | 127022.4 | -30.025 | <0.001 |
| Act | -1.394 | 0.179 | 136592 | -7.771 | <0.001 |
| Minute:r_t | 1.439 | 0.038 | 138516.1 | 37.74 | <0.001 |
| Minute:Act | 1.332 | 0.165 | 137993.9 | 8.076 | <0.001 |

**Table S2:** Model results table of linear mixed effects model for circular data, using the magnetic response for the entire test duration.

| **Predictor** | **Estimate** | **SE** | **df** | **t** | **p** |
|---|---|---|---|---|---|
| Intercept | 0.272 | 0.030 | 16.817 | 9.000 | <0.001 |
| Helper variable | 0.006 | 0.004 | 139366.988 | 1.711 | 0.087 |
| Minute | -0.372 | 0.038 | 7.484 | -9.738 | <0.001 |
| r_t | -0.382 | 0.028 | 97161.072 | -13.591 | <0.001 |
| Act | -0.479 | 0.179 | 126853.161 | -2.671 | 0.008 |
| Minute:r_t | 0.610 | 0.038 | 130274.539 | 16.002 | <0.001 |
| Minute:Act | 0.118 | 0.165 | 128386.126 | 0.717 | 0.474 |

**Table S3:** Model results table of linear mixed effects model for circular data, using the geographic trained response for the entire test duration.

| **Predictor** | **Estimate** | **SE** | **df** | **t** | **p** |
|---|---|---|---|---|---|
| Intercept | 0.053 | 0.039 | 13.422 | 1.349 | 0.2 |
| Helper variable | -0.075 | 0.004 | 139367 | -20.272 | <0.001 |
| Minute | -0.046 | 0.036 | 24.66 | -1.281 | 0.212 |
| r_t | -0.146 | 0.028 | 127807.2 | -5.159 | <0.001 |
| Act | 1.618 | 0.18 | 135820.9 | 8.997 | <0.001 |
| Minute:r_t | 0.208 | 0.038 | 137748.5 | 5.433 | <0.001 |
| Minute:Act | -1.946 | 0.165 | 134046.3 | -11.779 | <0.001 |

**Table S4:** Model results table of linear mixed effects model for circular data, using the trained magnetic response for the entire test duration.

| **Predictor** | **Estimate** | **SE** | **df** | **t** | **p** |
|---|---|---|---|---|---|
| Intercept | -0.118 | 0.035 | 14.648 | -3.351 | 0.004 |
| Helper variable | -0.054 | 0.004 | 139367 | -14.523 | <0.001 |
| Minute | 0.387 | 0.035 | 25.898 | 11.064 | <0.001 |
| r_t | 0.251 | 0.028 | 120794.7 | 8.864 | <0.001 |
| Act | -0.502 | 0.18 | 134242.1 | -2.785 | 0.005 |
| Minute:r_t | -0.677 | 0.038 | 134301.7 | -17.685 | <0.001 |
| Minute:Act | 0.991 | 0.166 | 132074.9 | 5.98 | <0.001 |

**Table S5:**
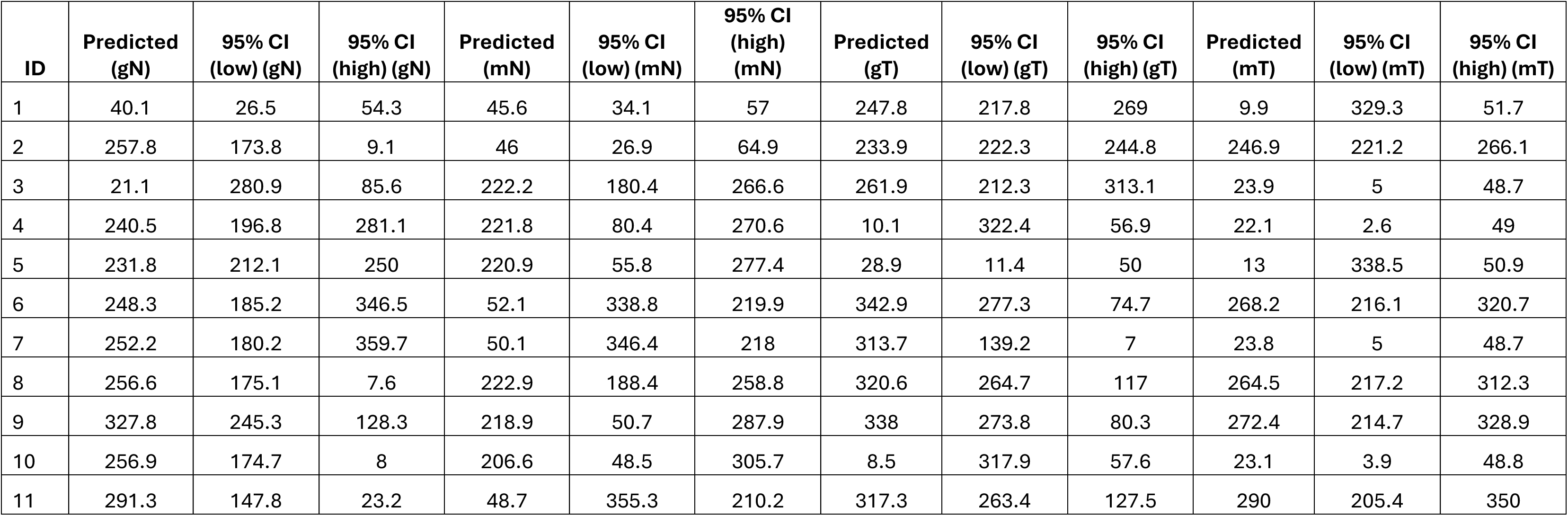
Individual model predictions corresponding to the individual results in Figure 2 (4^th^ column).

| **ID** | **Predicted (gN)** | **95% CI (low) (gN)** | **95% CI (high) (gN)** | **Predicted (mN)** | **95% CI (low) (mN)** | **95% CI (high) (mN)** | **Predicted (gT)** | **95% CI (low) (gT)** | **95% CI (high) (gT)** | **Predicted (mT)** | **95% CI (low) (mT)** | **95% CI (high) (mT)** |
|---|---|---|---|---|---|---|---|---|---|---|---|---|
| 1 | 40.1 | 26.5 | 54.3 | 45.6 | 34.1 | 57 | 247.8 | 217.8 | 269 | 9.9 | 329.3 | 51.7 |
| 2 | 257.8 | 173.8 | 9.1 | 46 | 26.9 | 64.9 | 233.9 | 222.3 | 244.8 | 246.9 | 221.2 | 266.1 |
| 3 | 21.1 | 280.9 | 85.6 | 222.2 | 180.4 | 266.6 | 261.9 | 212.3 | 313.1 | 23.9 | 5 | 48.7 |
| 4 | 240.5 | 196.8 | 281.1 | 221.8 | 80.4 | 270.6 | 10.1 | 322.4 | 56.9 | 22.1 | 2.6 | 49 |
| 5 | 231.8 | 212.1 | 250 | 220.9 | 55.8 | 277.4 | 28.9 | 11.4 | 50 | 13 | 338.5 | 50.9 |
| 6 | 248.3 | 185.2 | 346.5 | 52.1 | 338.8 | 219.9 | 342.9 | 277.3 | 74.7 | 268.2 | 216.1 | 320.7 |
| 7 | 252.2 | 180.2 | 359.7 | 50.1 | 346.4 | 218 | 313.7 | 139.2 | 7 | 23.8 | 5 | 48.7 |
| 8 | 256.6 | 175.1 | 7.6 | 222.9 | 188.4 | 258.8 | 320.6 | 264.7 | 117 | 264.5 | 217.2 | 312.3 |
| 9 | 327.8 | 245.3 | 128.3 | 218.9 | 50.7 | 287.9 | 338 | 273.8 | 80.3 | 272.4 | 214.7 | 328.9 |
| 10 | 256.9 | 174.7 | 8 | 206.6 | 48.5 | 305.7 | 8.5 | 317.9 | 57.6 | 23.1 | 3.9 | 48.8 |
| 11 | 291.3 | 147.8 | 23.2 | 48.7 | 355.3 | 210.2 | 317.3 | 263.4 | 127.5 | 290 | 205.4 | 350 |

**Figure S4:**
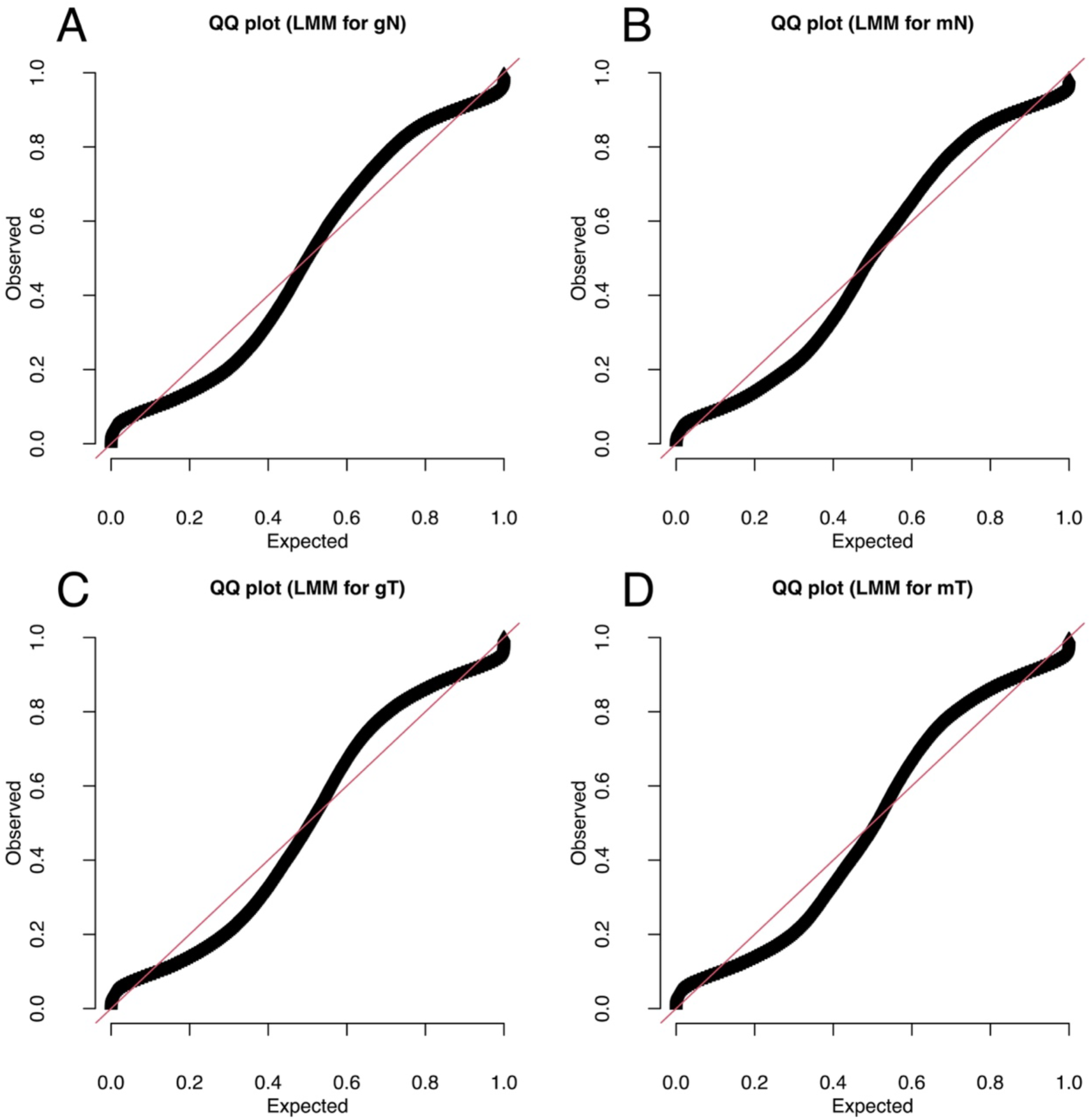
QQ plots for each of the four models used (A: towards geographic North, B: towards magnetic North, C: towards the geographic trained direction and D: towards the magnetic trained direction). The samples are clearly not normally distributed, as they are circular samples that have been linearised using their x and y components of the angles in radians, however, earlier work [10] has shown that this can be used in linear models. The slight S-shape of the (linear) residuals is likely due to the finite limits of the cosine and sine values (they can only be between -1 and 1).

**Figure S5:**
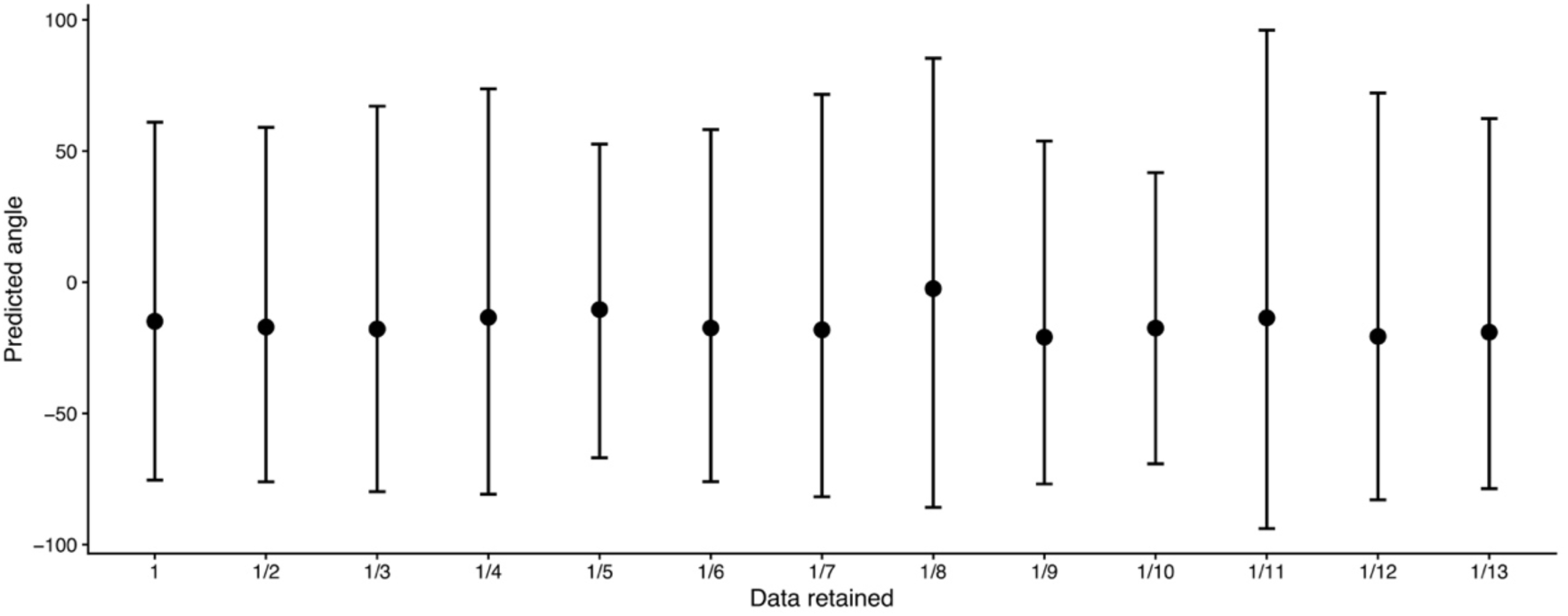
Model sensitivity analysis using data thinning for the magnetically trained response (mT). In this analysis we used the entire data set (indicated with ‘1’) followed by every second data point (‘1/2’), every third (‘1/3’) and so forth. At ‘1/14’ of the entire data set the model did not converge, hence we stopped there. The model used was the same as for the main analysis throughout. The predicted mean direction and the 95% confidence interval were similar throughout (note: ‘0’ is the expected direction, anticlockwise to this reference is indicated by negative and clockwise by positive numbers, all in degrees).

**Table S6:**
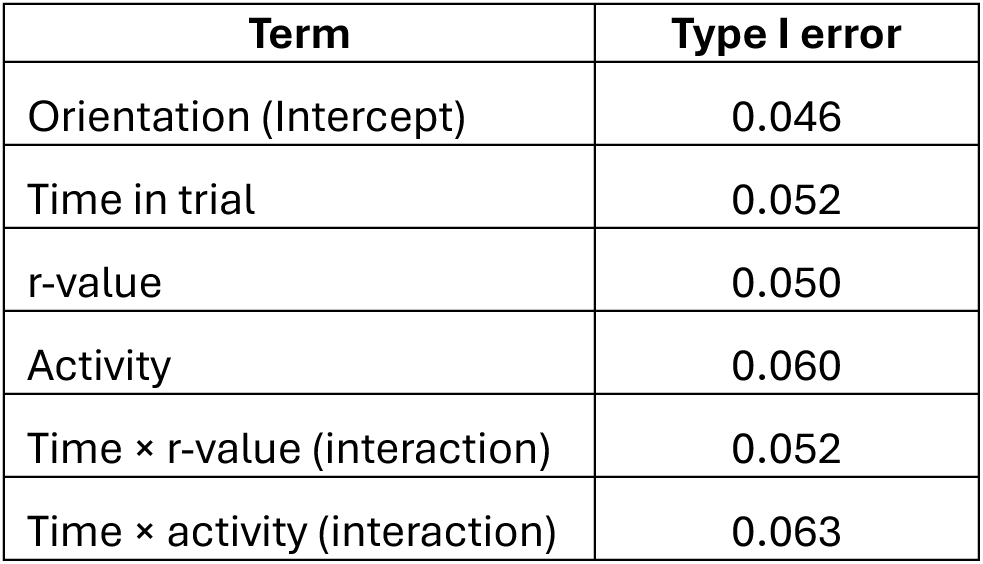
Type I error for each term used in the simulations of the LMM model. Values are the proportion of 1000 simulated uniform samples (fitted with the full model) in which the p-value of the respective term was below 0.05; values between 0.036 and 0.064 are expected by chance.

**Figure S6:**
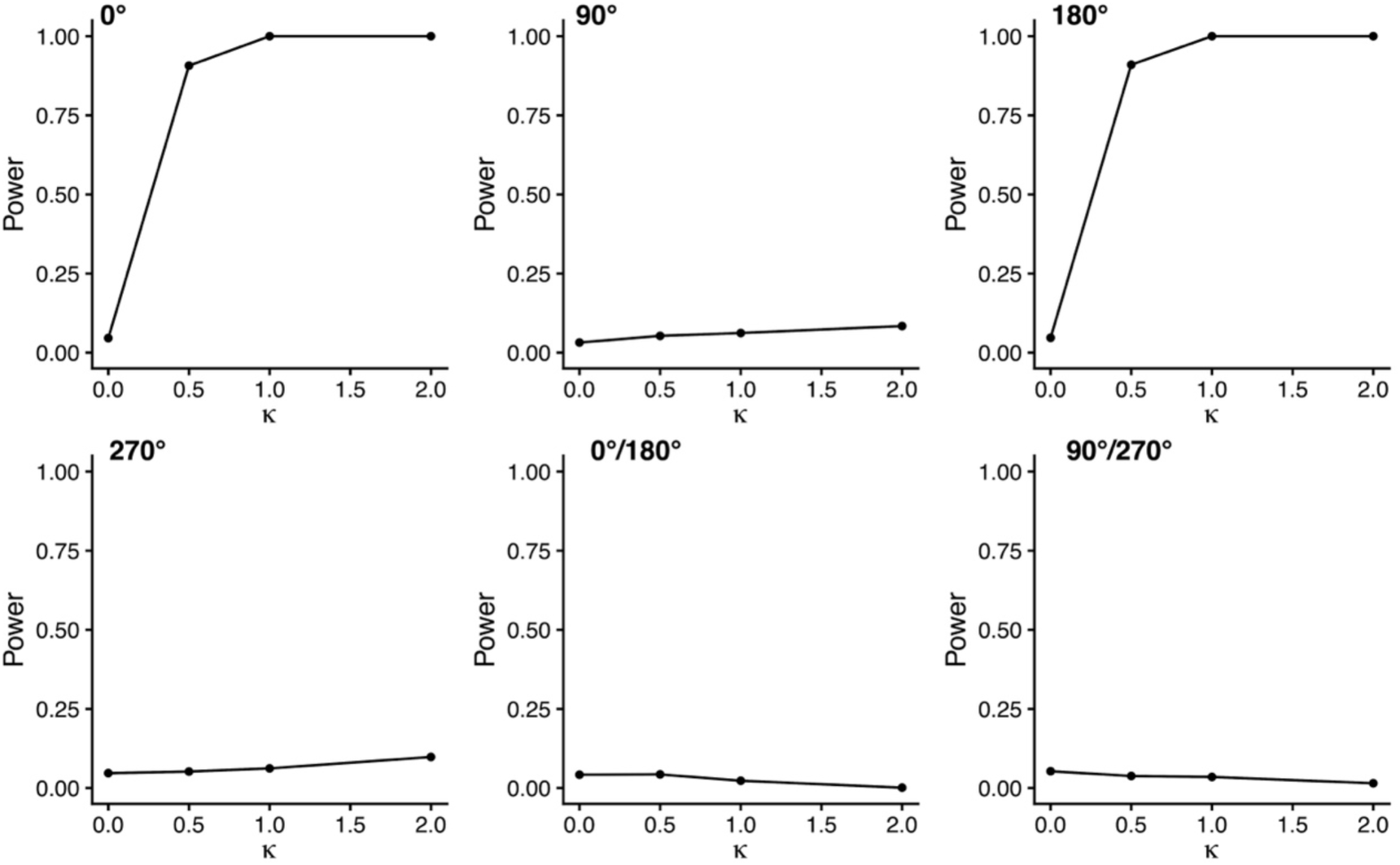
Power analysis results of intercept only model, when considering the p-value of the intercept, with cos(θ) as the base level of the helper variable. Testing the power to detect directionality of a von Mises sample based on direction (mean direction shown in each panel: unimodal: 0°, 90°, 180°, 270° and bimodal: 0°/180° and 90°/270°) and κ (ranging from 0 to 2). Power was only usable for clustering towards or opposite to 0°.

**Figure S7:**
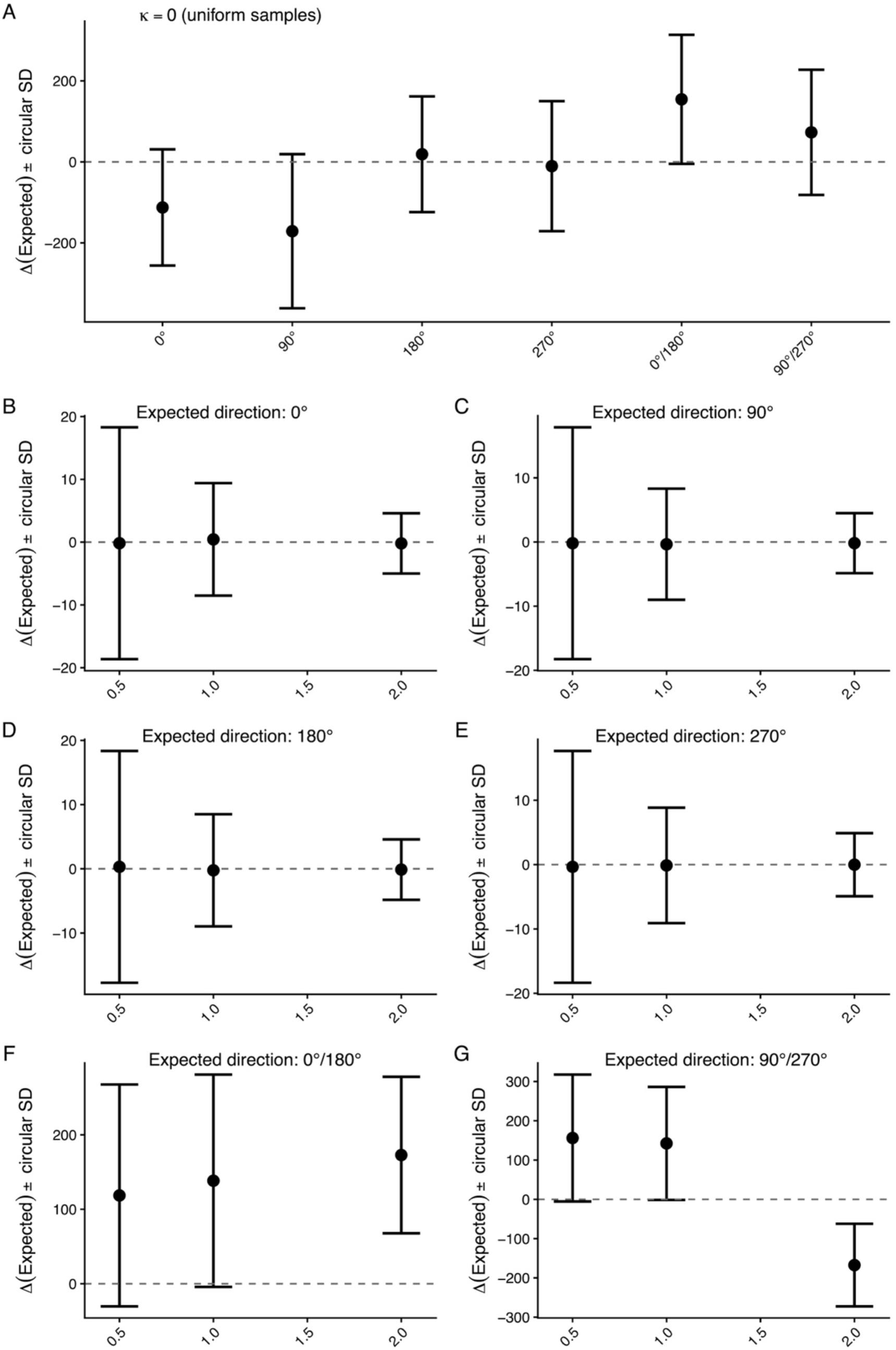
Simulation results of intercept only models. Random samples (A) resulted in highly variable and unusable mean predictions (circular standard deviation: circular SD). In contrast, samples from unimodal von Mises distributions with mean directions 0°, 90°, 180° and 270° had correct mean direction predictions with low circular standard deviations (note, y-axes differ between panels). However, bimodal (axial) samples resulted again in unstable mean predictions, diverted away from the correct direction.

**Figure S8:**
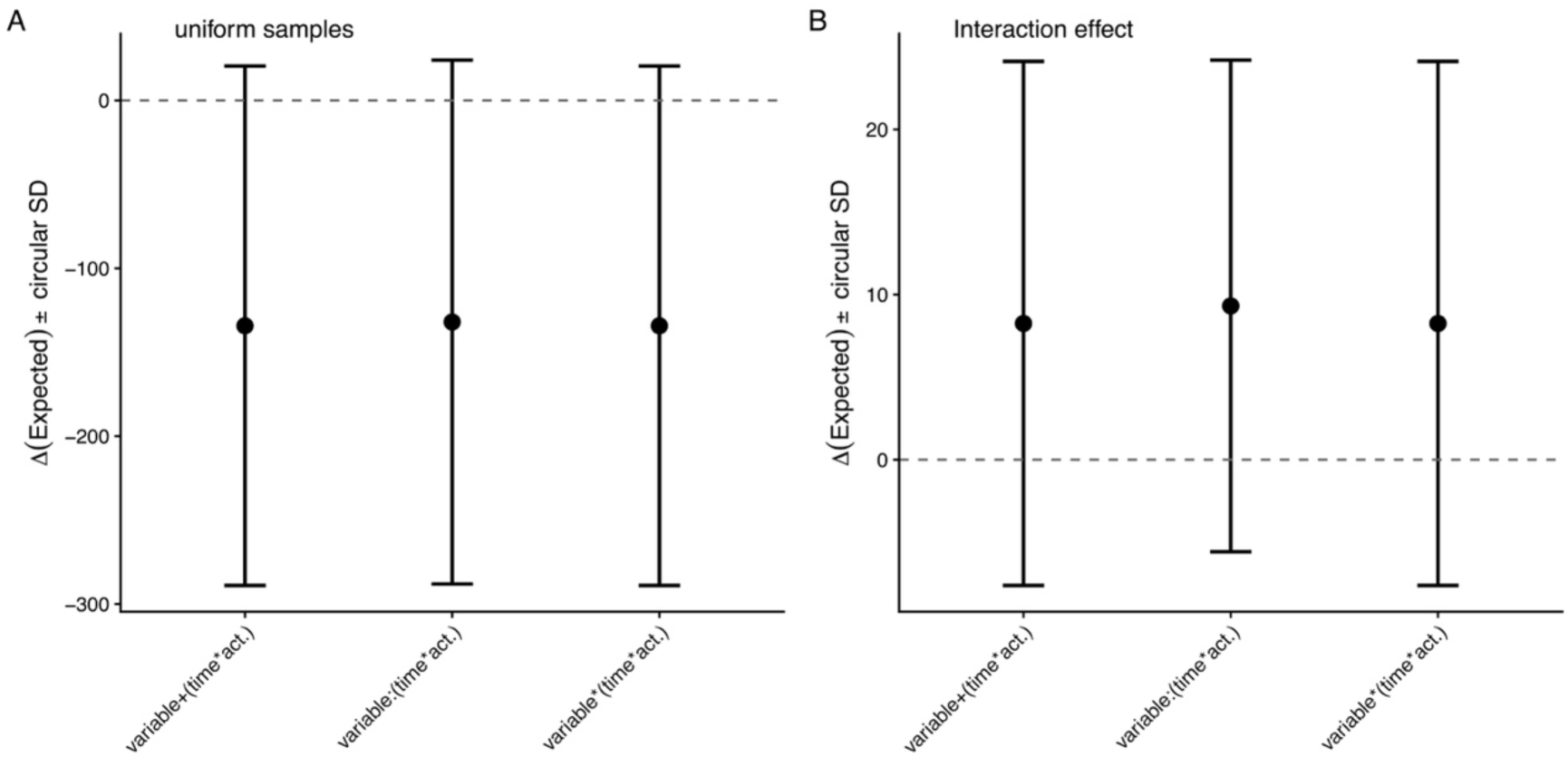
Simulation results of a more complex model, including time and activity (act), with the helper variable (variable) either as a main effect, only as an interaction or as a main and interaction effect with time and activity. The uniform samples showed highly variable unusable predictions for all models (A), all models stayed at nominal type I error levels when considering the p-value of the intercept (0.047, 0.047 and 0.043, respectively). All three approaches predicted the correct direction well (mean deviation 8-9°, circular SD 15-16°) and included the correct direction within their circular standard deviation. The slight positive deviation can be explained by the way the samples were simulated: increasing activity was associated with increasing tendency towards 180° and increasing time was associated with increasing concentration, as we simulated a positive interaction. Samples drawn from high activity and late in the trial were therefore highly clustered towards 180°, whereas samples from low activity and early in the trial resulted in less clustered orientation towards 0°. Because 90° was defined as the expected direction, the stronger clustering towards 180° biased the predictions towards values slightly above 90°. The power for the intercept p-value to detect deviation from uniformity was unusable throughout (0.069, 0.159 and 0.137, respectively), highlighting the issue of this approach with clustering towards 90° and bimodality in the data set.

**Figure S9:**
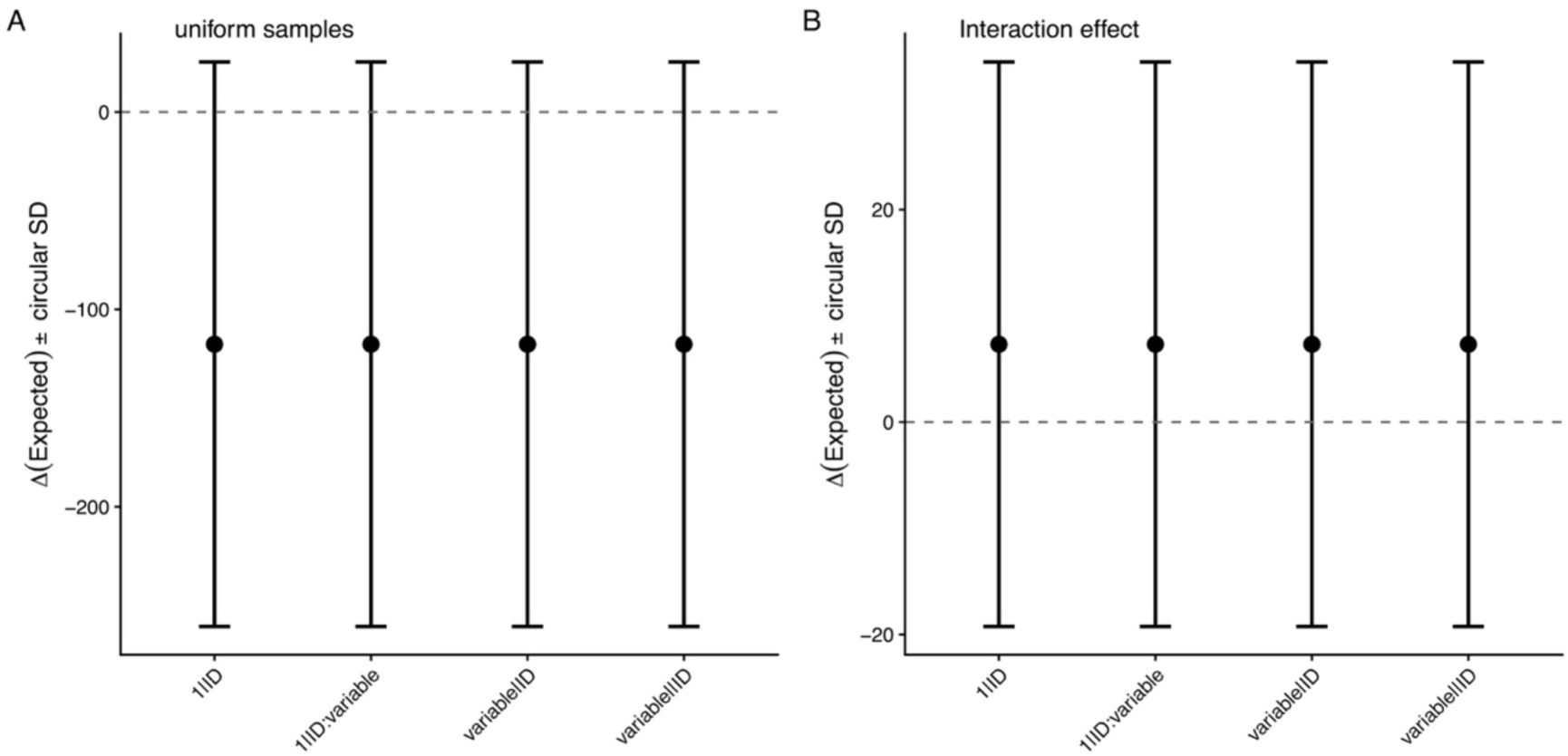
Simulation results of the more complex model adding repeated measures on the same ‘individuals’ and including this structure in a mixed effects model. The main effects were kept identical (formula: variable * (time * act)), however the random error term was either kept to the minimum (1 | ID), or the helper variable was added in one of the two following ways: (1 | ID : variable), (variable | ID) or (variable | |ID). Uniform samples produced highly variable and unusable predictions (A). However, predictions were acceptable for non-random samples (B) and the expected direction fell within the circular standard deviations of the predictions. In contrast to the results shown in Fig. S8 (above), the variation around the mean increased (circular SD 27° compared with 15-16°), but the directionality of the predictions stayed similar with a slight (and explainable) bias towards 180° (see explanation in Fig. S8). Predictions were identical between the four model structures tested. When considering the p-value of the intercept type I error stayed at nominal levels throughout (0.044, 0.044, 0.045, 0.045, respectively); and power was unusably low for non-random samples (0.099, 0.091, 0.091 and 0.091, respectively).

